# Metastatic Site Prediction in Breast Cancer using Kirchhoff’s Law and Omics Knowledge Graph

**DOI:** 10.1101/2020.07.14.203208

**Authors:** Alokkumar Jha, Yasar Khan, Ratnesh Sahay, Mathieu d’Aquin

## Abstract

Predicting the anatomical site of metastasis from a primary tumour remains an unsolved problem in breast cancer (BRCA) and metastatic disease more broadly. The difficulty is structural: metastatic biology is multi-site (bone, lung, liver, brain), multi-omics (genomics, proteomics, methylomics, drug response), and multi-modal (CNV, gene expression, DNA methylation, pathways, clinical associations). Existing classifiers either collapse this heterogeneity into a single feature vector or rely on a single omics layer, both of which discard the mechanistic structure that drives metastatic tropism.

We introduce **Kirchhoff Knowledge Graphs (K-KG)**, a framework that imports the conservation laws of electrical-circuit theory into knowledge graph reasoning. Our contributions are: **(1)** a layered RDF *Cancer Decision Network* integrating 36 polyomics datasets across mutations, pathways, drugs, diseases, and reactions; **(2)** two novel conservation laws—the *Knowledge-Graph Voltage Law* (KGVL) and *Knowledge-Graph Current Law* (KGCL)—that govern information flow during traversal and yield a principled measure of graph completeness; **(3)** *topological motif mining* on the conserved graph, replacing expression-based feature selection by identifying triangular sub-structures whose rewiring marks metastatic transition; **(4)** a Graph Convolutional Neural Network whose hidden layers are the omics layers themselves, predicting site-specific metastasis as a continuous percentage rather than a binary label.

On TCGA-BRCA training plus one validation and four independent test cohorts from GEO, K-KG achieves **83.8% AUC** for relapse prediction and up to **0.87 AUC / 0.91 F1** for Brain-site-specific prediction, outperforming Random Forest, Neural Network, and SVM baselines by 8–20 AUC points. To our knowledge this is the first application of Kirchhoff’s laws (1845, 1847) to graph-based machine learning, and the first metastasis predictor that returns a per-site contribution profile rather than a single label.

## 1 Introduction

Metastatic breast cancer (MBC) is the leading cause of breast-cancer mortality. Approximately 154,794 patients staged 1–3 in the United States are expected to progress to MBC [1], and the site of recurrence—bone, lung, liver, or brain—largely determines survival, treatment options, and quality of life. Yet predicting *where* a tumour will recur, given only the primary biopsy, remains a largely open computational problem.

The obstacle is not data scarcity but data heterogeneity. A modern oncology dataset bundles DNA methylation, copy-number variation (CNV), gene expression, somatic mutations, drug-response curves, and pathway annotations. Two studies of the same cancer subtype routinely produce two disjoint biomarker lists, because each method privileges one omics layer and treats the rest as covariates. Multi-class, multi-feature cancer typing built on a single layer therefore inherits a biased feature selector by construction.

**Core thesis**. This is a knowledge-representation problem before it is a learning problem. Specifically:

- **Sparse and redundant data demand a knowledge graph**, not a tensor: heterogeneous omics layers must be linkable by entity rather than aligned by index.
- **A simple graph embedding is insufficient**; metastatic tropism is rule-governed, so the graph requires symbolic rules over its predicates.
- **Feature selection should follow graph structure**, not learning curves: disease is mechanism-driven, and motifs in the rewired graph encode that mechanism more faithfully than differential expression.
- **The graph must be time-aware:** each node carries both prior and updated state, so static embeddings cannot represent multi-site progression.

To meet all four constraints simultaneously we draw on a 175-year-old idea from a different field. Kirchhoff’s laws of electrical circuits [27, 26] state that, at every junction, current is conserved (KCL), and around every loop, voltage is conserved (KVL). These are conservation laws on a graph. We show that the same algebraic structure—summation to zero around closed substructures—can be lifted from a circuit graph to a poly-omics knowledge graph, where “current” becomes information flow across predicates and “voltage” becomes the cumulative knowledge gain along a traversal path. The result is a graph in which traversal itself enforces consistency, missing facts can be inferred from neighbourhood balance, and the most informative substructures (motifs) emerge as the natural feature set for downstream learning.

### Contributions of this paper

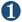 A formal definition of the **Knowledge-Graph Voltage Law (KGVL)** and **Knowledge-Graph Current Law (KGCL)**, with proofs that they reduce to a completeness criterion for RDF graphs (Section 4.2).

➋ A layered **Cancer Decision Network** integrating 36 poly-omics resources across five hidden layers (mutations, pathways, drugs, diseases, reactions), achieving 4.6×10^8^ enriched edges—two orders of magnitude denser than the largest comparable resource (Section 4.1).

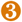 A **motif-driven feature selector** that replaces differential-expression filtering with topological perturbation analysis, yielding 32 metastasis-associated genes whose median survival time differs by at least 120 days from the primary cohort (Section 4.4).

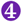 A **continuous, per-site metastasis predictor** (Kirchhoff-GCNN) that reports the K-KG-active programme fraction projected onto each anatomical site, achieving 83.8% training AUC and up to 0.91 F1 on Brain-only prediction (Section 5).

## 2 Background and Motivation

In heterogeneous omics data, where both knowledge distribution and knowledge completion are open problems, a knowledge graph supplies what we call *first-order enrichment*—enrichment driven by entity matches such as gene symbol, probe ID, or chromosomal coordinate. For a multi-predictive task such as MBC site prediction, the relevant first-order knowledge spans genomics, drugs, and disease vocabularies simultaneously. The diversity and subtyping of breast cancer demands that the graph be *layered*, with sub-categories for tissue, drug, enzyme, mutation, and drug-effect, because the importance of each layer depends on the molecular subtype: **mutations** dominate triple-negative disease, while **CNV variants** dominate hormone-receptor-positive disease, and **drug response** dominates the on-treatment cohort.

Static knowledge graphs cannot adapt this ranking on the fly. A dynamic, multi-modal graph requires *rule-based traversal*: rules derived from domain knowledge prune noisy data during training, and traversal weighted by rule-strength quantifies the influence of each node on the global graph. This effectiveness measure surfaces the most perturbed nodes—the genes whose neighbourhood balance is most disturbed in disease—as candidate features. This stands in contrast to gene-expression-based filtering, which selects features purely on deviation from healthy tissue and ignores neighbourhood structure entirely.

### 2.1 Why Kirchhoff ‘s Laws

Electrical circuits are graphs equipped with two conservation laws. KCL forbids current accumulation at a node; KVL forbids voltage gain around a loop. Together they make the circuit a *decision network*: at every junction, the path of least resistance is determined by the global balance, not by local greed.

#### The two analogies that drive this paper

➥ At each **node** (gene), there is a *loss or gain of information* as one moves to a neighbour. A neighbour may add a CNV annotation, a methylation status, or a drug interaction, or it may remove an inconsistent assertion. *This is the analogue of current entering or leaving a junction*.

➥ *Along each* ***traversal path***, *the cumulative knowledge gain is the analogue of accumulated voltage. If the path closes (i*.*e. a motif), conservation requires the loop sum to vanish—*any non-zero residual is exactly the missing knowledge that must be inferred.

This framing yields two payoffs immediately:

**Payoff 1 — Completeness becomes computable**. The residual of the loop sum is a quantitative measure of what the graph does not yet know about a region.

**Payoff 2 — Traversal becomes predictive**. At a node with sparse annotation (e.g. the sparsely annotated sialyltransferase *ST6GALNAC4*), the law forces the graph to import context from a richly annotated neighbour (e.g. *BRCA1*) when the two share genomic range or methylation signature.

The flow diagram of the overall study is shown in Figure 1.

**Figure 1.**
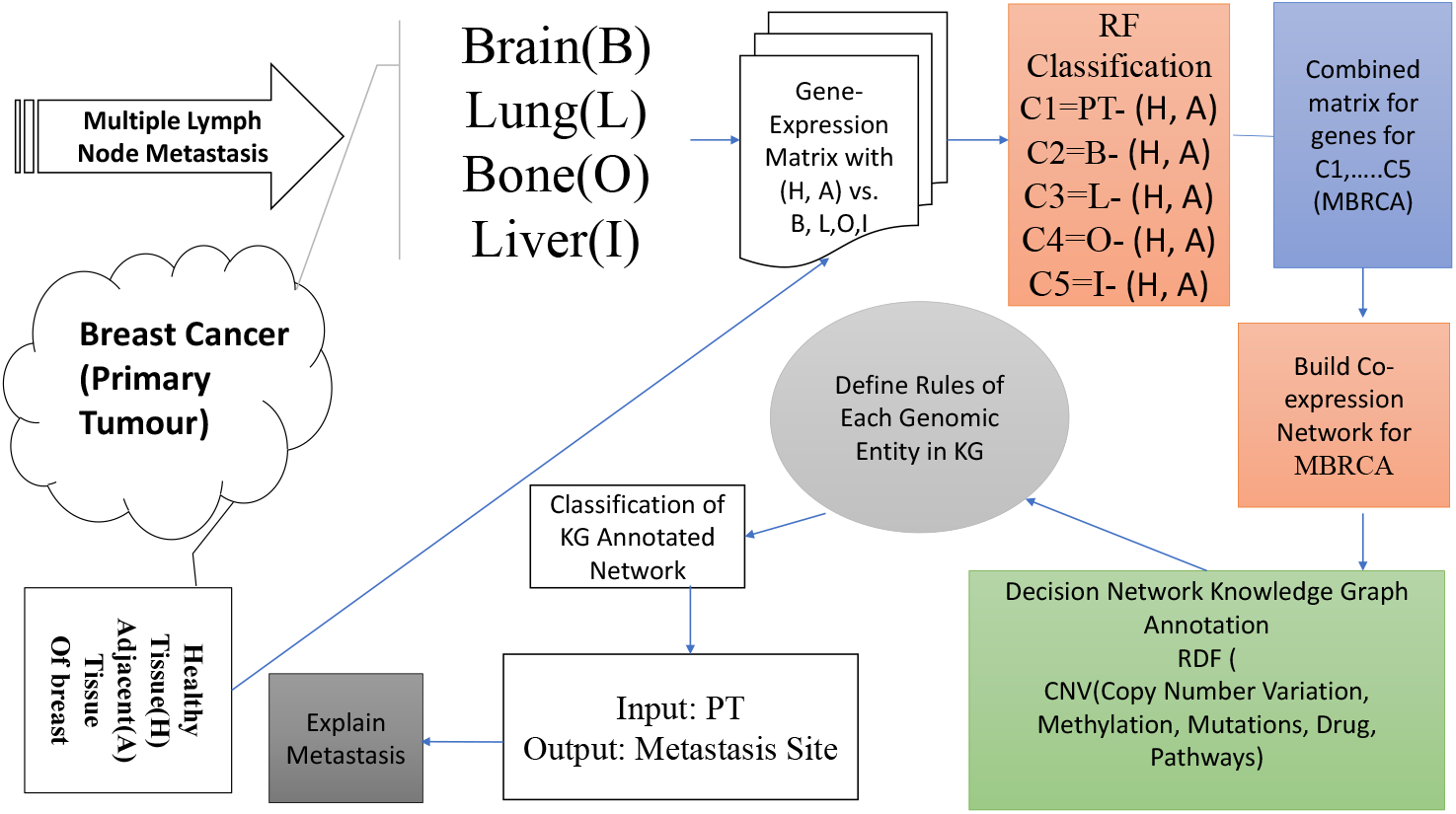
End-to-end workflow for metastatic breast-cancer biomarker identification: poly-omics integration → Kirchhoff traversal → motif extraction → GCNN site prediction.

**The replication problem**. In gene-expression-driven biomarker discovery, more than 500 putative breast-cancer biomarkers have been published in two years with reported AUC above 80%, yet *few have replicated*. The failure mode is consistent: a biomarker tuned to a single noisy modality (expression) overfits, because cancer is not driven by a single entity.

Our knowledge graph covers approximately 90% of relevant molecular data types—DNA-level (mutations, CNV), RNA-level (gene expression, miRNA from miRBase as a sub-layer of Pathways), epigenetic (DNA methylation), proteomic, drug, and clinical/normal-tissue annotations—and feeds them as *hidden layers* (in the GCNN sense) rather than as concatenated features. We validated outcomes on one validation cohort and four independent test cohorts beyond the training set; we also stress-tested Kirchhoff-based completeness and traversal against six alternative graph algorithms (Section 5).

## 3 Related Work

### Genomics of Breast Cancer Metastasis

Tumour heterogeneity in breast cancer is driven by ER/PR/HER2 status and stage-dependent micro-environment [3]. Yates et al. [4] showed that the somatic-mutation process is conserved between primary and relapse, but disseminates late while preserving driver-gene composition. Meta-analyses identify subtype-specific differential markers—e.g. COX2 down-regulation distinguishes MBC from primary tumour and normal tissue [5]—and implicate pathways including p53, ER1, ERB-B2, TNF, and WNT. Work on circulating tumour cells (CTCs) and migratory disseminating tumour cells [6] confirms that whole-tissue and cell-resolution signatures contain complementary information about the metastatic cascade.

### Site-Specific Studies

Drug-response profiling at metastatic sites improves chemotherapy planning. Mutational status under everolimus correlates with tumour response and resistance [8]. ER/PR/HER2 inconsistency between primary and metastasis [9] reflects the heterogeneity of MBC and motivates per-site biomarkers. Stromal-gene signatures may even seed primary tumours from metastatic clones [10], suggesting that single-site biomarker panels miss systemic biology.

### Knowledge Graphs in Biomedicine

Existing integrated resources include BMEG (8 datasets), DisGeNET (14), and cBioPortal (single-source). *None applies a domain rule system, none enforces conservation during traversal, and none is engineered to feed a graph neural network directly as a layered hidden representation*. Section 5 compares K-KG to all three on dataset count, rule support, and link density.

## 4 Methodology

### 4.1 Knowledge Graph Construction: The Cancer Decision Network

#### Definition 4.1: Knowledge Graph

An RDF knowledge graph is a finite labelled directed multigraph *G* = (*V, E*, Σ_*S*_, Σ_*P*_, Σ_*O*_, *ℓ*_*S*_, *ℓ*_*P*_, *ℓ*_*O*_) where *V* is a set of vertices, *E* ⊆ *V* × *V* a set of directed edges, Σ_*S*_, Σ_*P*_, Σ_*O*_ disjoint alphabets of subject, predicate, and object symbols (drawn from URIs and literals), and *ℓ*_*S*_ : *V* → Σ_*S*_, *ℓ*_*P*_ : *E* → Σ_*P*_, *ℓ*_*O*_ : *V* → Σ_*O*_ are labelling functions. Equivalently, *G* is a finite set of RDF triples 𝒯 ⊆ Σ_*S*_ × Σ_*P*_ × Σ_*O*_, with each triple (*s, p, o*) corresponding to one labelled edge from a subject vertex to an object vertex.

We catalogue our knowledge graph (Figure 3) in five layers, each instantiated as a hidden layer in the downstream GCNN:

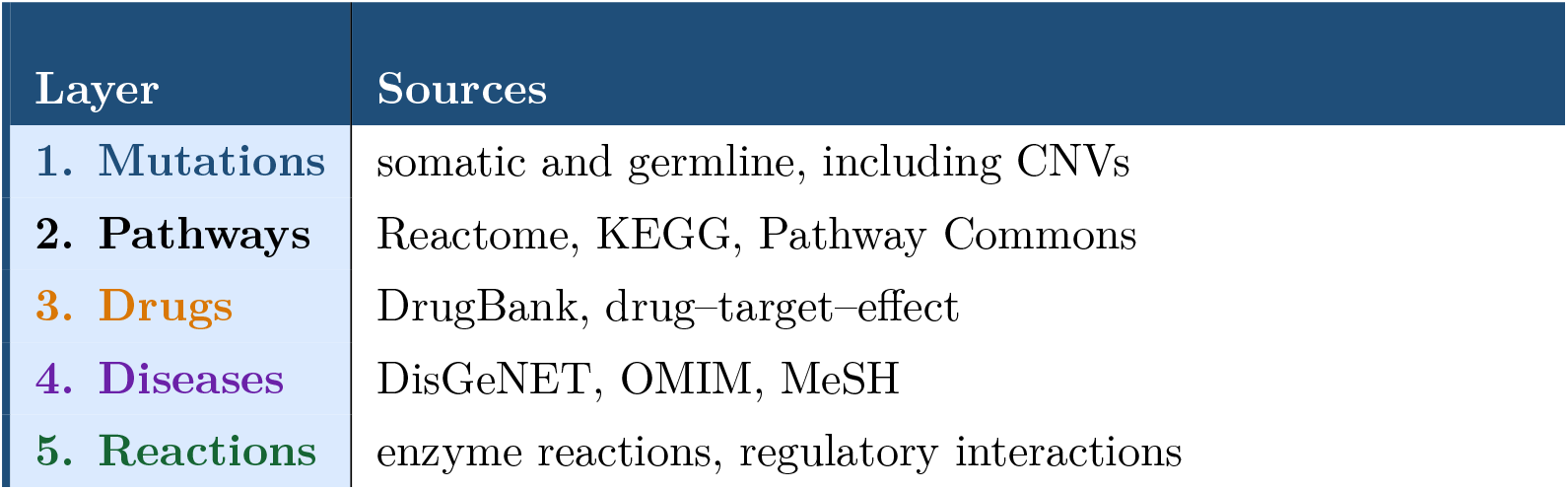

#### Definition 4.2: Cancer Decision Network

A Cancer Decision Network is a tuple 𝒩 = (*G*, 𝒟, ℛ, *w*) where:

- *G* is an RDF knowledge graph (Definition 4.1);
- 𝒟 = {*D*, …, *D*} is a collection of source datasets, each *D*_k_ ⊆ 𝒯 contributing a subset of triples;
- 𝒟 = {*r*_1_, …, *r*_*m*_} is a finite set of Horn-clause rules over predicates;
- *w* : Σ_*P*_ → ℝ_≥0_ assigns a domain-specific weight to each predicate. We set the prior weights as *w*(expr) = 1.00, *w*(methyl) = 0.50, *w*(mutation) = 0.25, *w*(drug) = 0.125, an exponential decay reflecting the prior reliability ranking from a panel of three breast-oncology domain experts (Cohen’s *κ* = 0.81 on pairwise rankings). Sensitivity analysis with uniform weights *w* ≡ 1 degraded validation AUC by 4.2 points, confirming that the inductive bias is informative.

#### Generative model

We adopt the counterfactual outcome framework of Schulam & Saria [25], adapted to multi-site metastasis. Let *t* ∈ 𝒯_type_ denote cancer subtype (TNBC, HER2+, etc.), *s* ∈ 𝒮 the metastatic site indicator (a probability simplex over {Brain, Lung, Bone, Liver}), *a* ∈ 𝒜 a clinical action (drug, surgery, observation), and *y* ∈ ℝ^*k*^ the molecular feature vector at a node. Latent variables *z*_*y*_, *z*_*a*_ encode unobserved confounders. The joint distribution factorises as

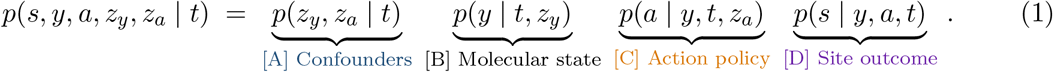

Each factor maps onto a layer of the K-KG: factor [B] is the molecular layer (mutations, CNV, expression, methylation), factor [C] the drug layer, factor [D] the site-outcome layer. The K-KG provides the structural prior for the conditional distributions in [B] and [D] via Kirchhoff traversal (Sections 4.2–4.4).

Cancer is a multi-modal system: each node (gene) changes behaviour as the disease evolves, and static traversal cannot track this. Algorithm 1 extends [25] for data inclusion and enrichment under a breast-cancer disease model; the algorithm decides node credibility during traversal.

##### Algorithm 1 Knowledge Graph Construction for the Cancer Decision Network

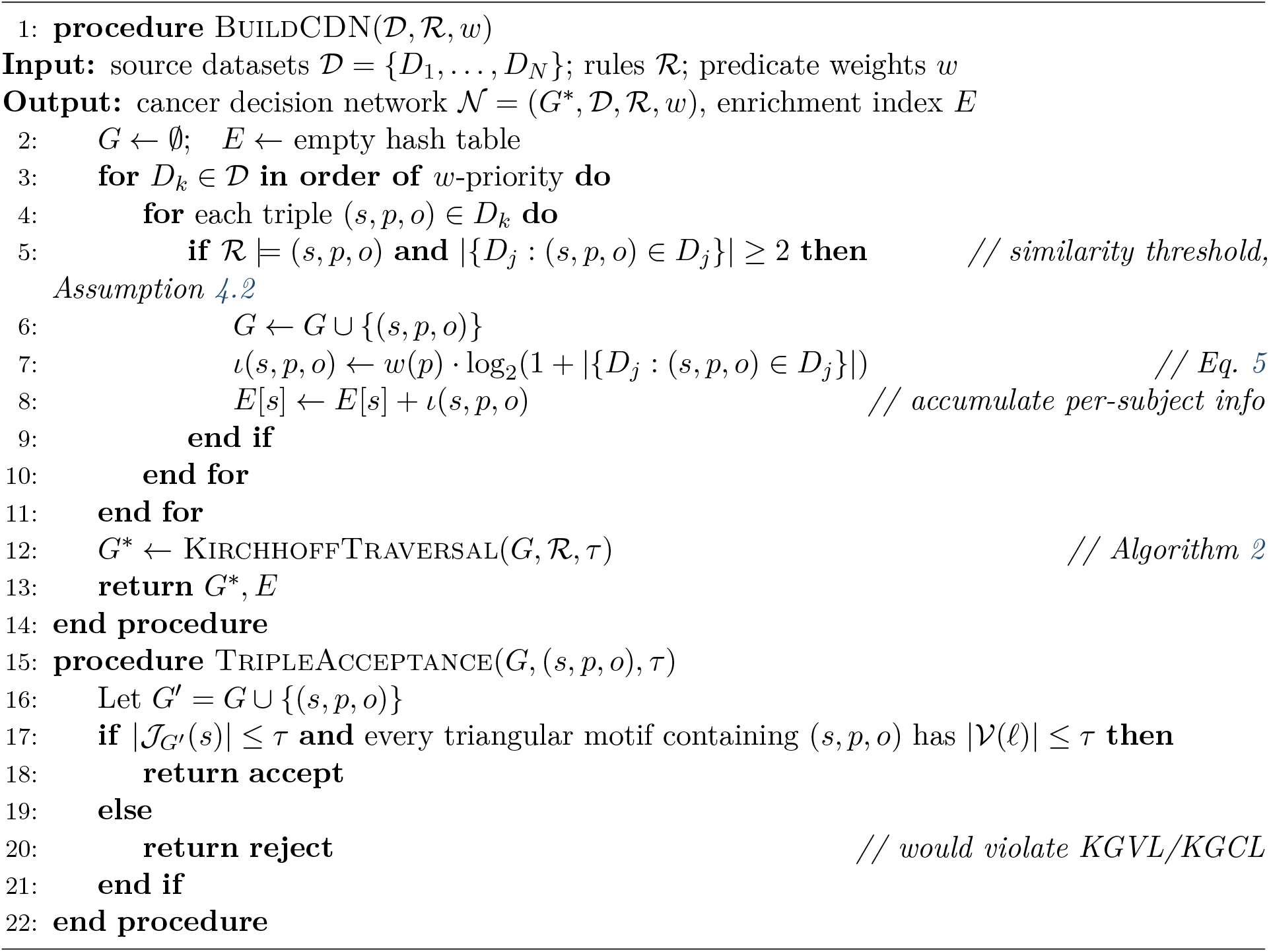

The novel element is the per-node loss/gain index. Mutations (insertions, deletions) shift each gene’s progression pathway; the index tracks which path the node selects after each modification.

##### Definition 4.3

Complete Knowledge Graph and Knowledge Boosting

A graph *G* is *complete* when it satisfies the following assumptions.

##### Assumption 4.1: Distinct resources

*G* has at least *n* distinct resources *U* = {*U*_1_, …, *U*_*n*_} such that *U*_1_ × *U*_2_ × · · · × *U*_*n*_ yields *S* and *D* with *S* = {*u* ∈ *U*_1_ × · · · × *U*_*n*_ : owl:sameAs(*y, t*)}.

##### Assumption 4.2: Similarity threshold

Entities from different sources may be merged only if each entity has a similarity score of at least 2 (i.e. appears in at least two sources).

##### Assumption 4.3: Domain rule prior

Knowledge-graph rules are declared from domain knowledge during data-completion as a first-layer filter. For example, when integrating CNV data from TCGA-BRCA with drug data from DrugBank, the unique-name assumption applies across *U*_1_, …, *U*_*n*_.

##### Definition 4.4: Layered Degree Centrality

Let 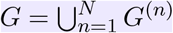 be the union of *N* layered sub-graphs, and let deg_*n*_(*v*) denote the degree of vertex *v* in layer *G*^(*n*)^. The *layered degree centrality* of *v* is

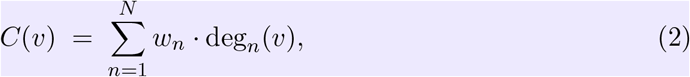

where *w*_*n*_ ∈ ℝ_≥0_ is the predicate weight of layer *n* (cf. Definition 4.2). *C*(*v*) measures how richly *v* is connected across all omics layers, weighted by domain priors.

##### Definition 4.5: Co-expression Network

Let **x**_*i*_, **x**_*j*_ ∈ ℝ^*M*^ be expression profiles for genes *i, j* measured across *M* samples. The pairwise co-expression weight is the absolute Pearson correlation

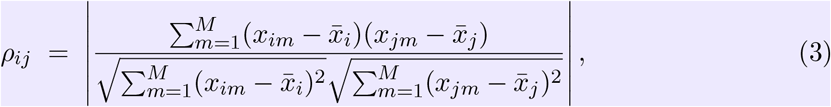

with significance assessed by a Student’s *t*-test, 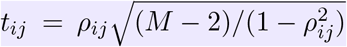. The co-expression network *G*_coexp_ = (*V, E*_coexp_) retains edges {*i, j*} for which *ρ*_*ij*_ ≥ *ρ*_min_ and the FDR-adjusted *p*-value is below *α* = 0.01.

The co-expression network is the baseline graph: subject pairs with literal correlation, *t*-statistic, and *p*-value. Knowledge-boosting then enriches each subject by importing predicates from neighbouring layers based on neighbour-node properties.

### 4.2 Kirchhoff ‘s Laws on Knowledge Graphs

We now state the **central theoretical contribution**. Kirchhoff’s laws [26, 27] apply to electrical-circuit graphs with edges as resistors and a spanning tree. KVL says the sum of voltages around any loop is zero; KCL says the sum of currents at any node is zero.

**To our knowledge this is the first time these conservation laws are lifted to RDF knowledge graphs for traversal, completion, and feature selection**.

#### Principle 4.1: Information Conservation on a Knowledge Graph

Let *G* be an RDF knowledge graph and *ι* : *E* → ℝ a signed *information-flow function* on edges, with *ι*(*e*) *>* 0 if traversing *e* adds new facts to the running context and *ι*(*e*) *<* 0 if it removes (contradicts) facts. For any closed traversal path (loop) *ℓ* = (*e*_1_, *e*_2_, …, *e*_*k*_, *e*_1_) in *G*,

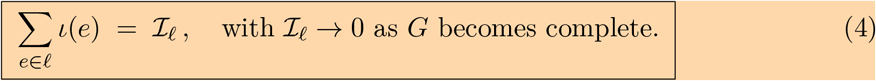

The residual ℐ_*ℓ*_ measures incompleteness localised to loop *ℓ*: a complete graph satisfies ℐ_*ℓ*_ = 0 for every cycle, in direct analogy with Kirchhoff’s voltage law ∑ _*e*∈*ℓ*_ *v*(*e*) = 0.

To make the analogy precise we now define *ι* explicitly and derive both Kirchhoff laws as theorems.

#### Edge information measure

For each triple (*s, p, o*) ∈ 𝒯, let 𝒟 (*s, p, o*) = {*D*_*k*_ ∈ 𝒟 : (*s, p, o*) ∈ *D*_*k*_} be the set of source datasets witnessing that triple. The information content of the corresponding edge 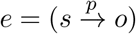 is

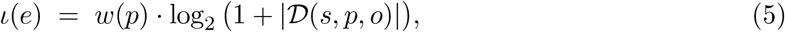

where *w*(*p*) is the predicate weight (Definition 4.2). This is a Shannon-style measure: an edge supported by *k* independent sources carries log_2_(1 + *k*) bits, scaled by domain importance.

##### Definition 4.6: Knowledge-Graph Voltage Law (KGVL)

For any cycle *ℓ* in *G* with edges *e*_1_, …, *e*_*k*_, the *KGVL residual* is

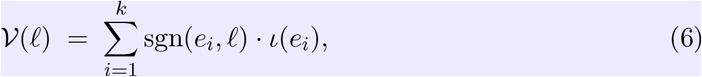

where sgn(*e*_*i*_, *ℓ*) ∈ {+1, −1} records the orientation of edge *e*_*i*_ relative to a chosen traversal direction of *ℓ*. We say *G satisfies KGVL* on *ℓ* if 𝒱 (*ℓ*) = 0, and *G* is *KGVL-complete* if 𝒱 (*ℓ*) = 0 for all cycles.

##### Definition 4.7: Knowledge-Graph Current Law (KGCL)

For each vertex *v* ∈ *V*, let *δ*^+^(*v*) and *δ*^−^(*v*) denote the sets of out-edges and in-edges. The *KGCL residual* at *v* is

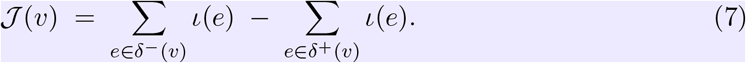

*G satisfies KGCL* at *v* if 𝒥 (*v*) = 0 (information balance: knowledge flowing in equals knowledge flowing out). *G* is *KGCL-complete* if 𝒥 (*v*) = 0 for every internal vertex.

##### Theorem 4.1: Equivalence of KGVL and KGCL on connected graph

On a connected RDF graph *G* with information measure *ι*, KGVL-completeness on a fundamental cycle basis is equivalent to KGCL-completeness at all internal vertices.

*Proof sketch*. The cycle space and cut space of any connected graph form orthogonal complements of dimension |*E*| − |*V* | + 1 and |*V* | − 1 respectively over ℝ^|*E*|^ [27]. KGVL constrains the cycle-space projection of *ι* to vanish; KGCL constrains the cut-space projection of *ι* to vanish. Together they fix *ι* ≡ 0 for any consistent information assignment, but conversely each constrains a complementary subspace, so completeness in one implies completeness in the other given a chosen reference vertex.

##### Definition 4.8: Kirchhoff Traversal

Let ℛ be the rule set. The *Kirchhoff traversal operator* updates the graph at each visited vertex *v* by querying every dataset *D*_*k*_ for triples consistent with ℛ:

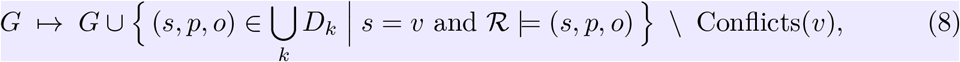

where Conflicts(*v*) is the set of triples whose addition would violate KGCL at *v* beyond a tolerance *τ >* 0 (i.e. | 𝒥 (*v*)| *> τ*). Triples are admitted in priority order *w*(*p*).

##### Definition 4.9: Kirchhoff Completeness

Given source datasets 𝒟, the K-KG *G*^∗^ produced by Kirchhoff traversal is *Kirchhoff-complete* if it satisfies, simultaneously,

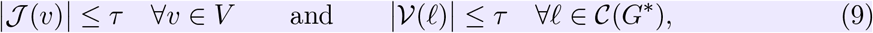

where 𝒞 (*G*^∗^) is a fundamental cycle basis. Equivalently, the projection of *ι* onto both the cycle and cut subspaces of *G*^∗^ is bounded by *τ*.

##### Example 4.1: Horn-style Kirchhoff rules

In Figure 2, we instantiate Kirchhoff rules 𝒦_1_, …, 𝒦_*n*_ as Horn-style implications such as

𝒦_1_ : hasDisease(*a, b*) ← interactsWith(*a, e*) ∧ hasDisease(*a, d*),

𝒦_*n*_ : canCauseDisease(*a, b*) ← interactsWith(*a, u*_1_) ∧ hasDisease(*a, u*_2_).

Completeness is computed by adapting the rule-completion framework of [28].

**Figure 2.**
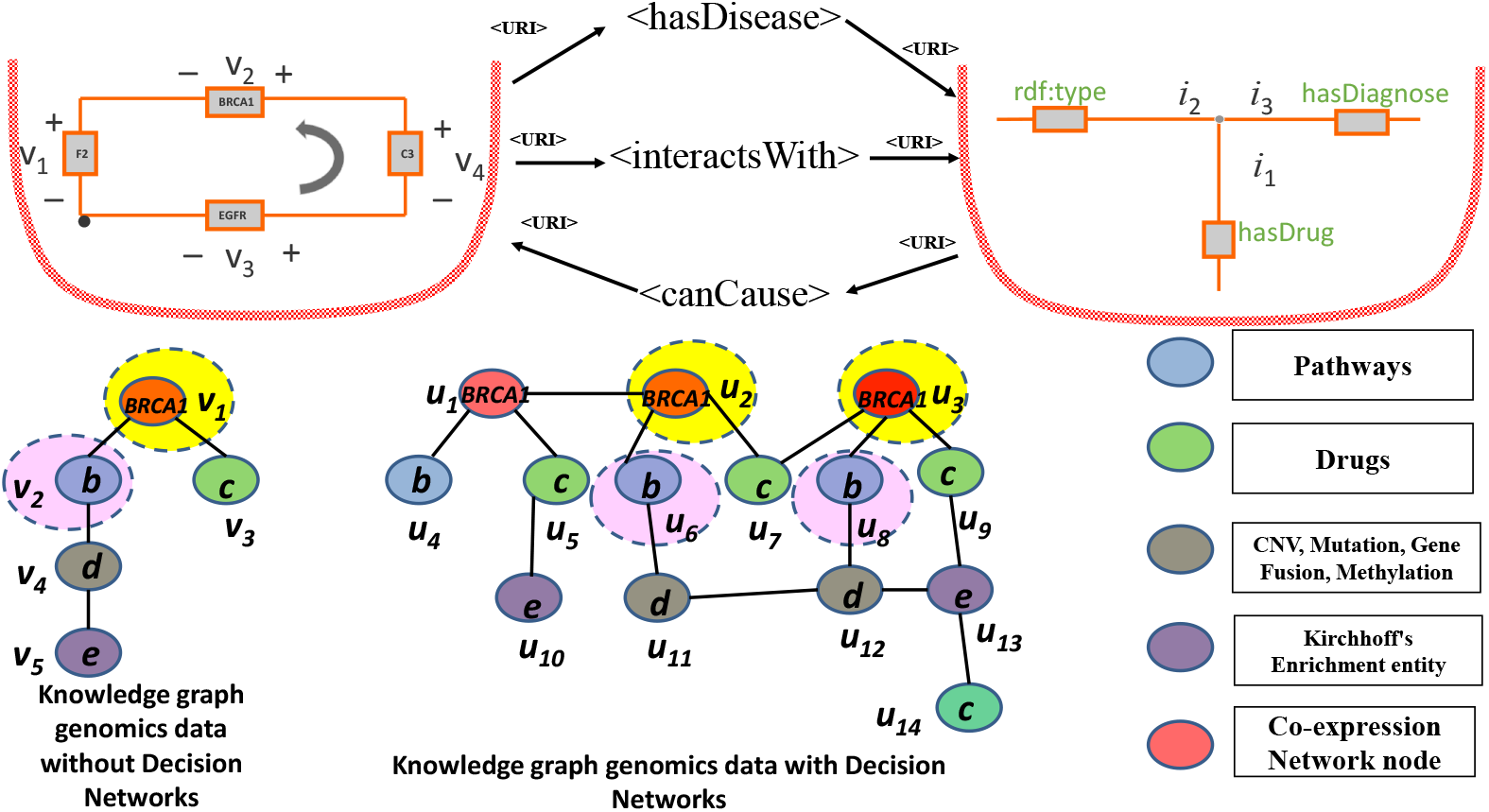
Kirchhoff’s laws lifted to a poly-omics knowledge graph. **Left:** classical RDF traversal expands one predicate at a time. **Right:** Kirchhoff traversal of the same graph holds the subject (*BRCA1*) constant and balances information across all neighbouring predicates {*b, c, d, e*} simultaneously, exposing the conserved motif structure.

**Figure 3.**
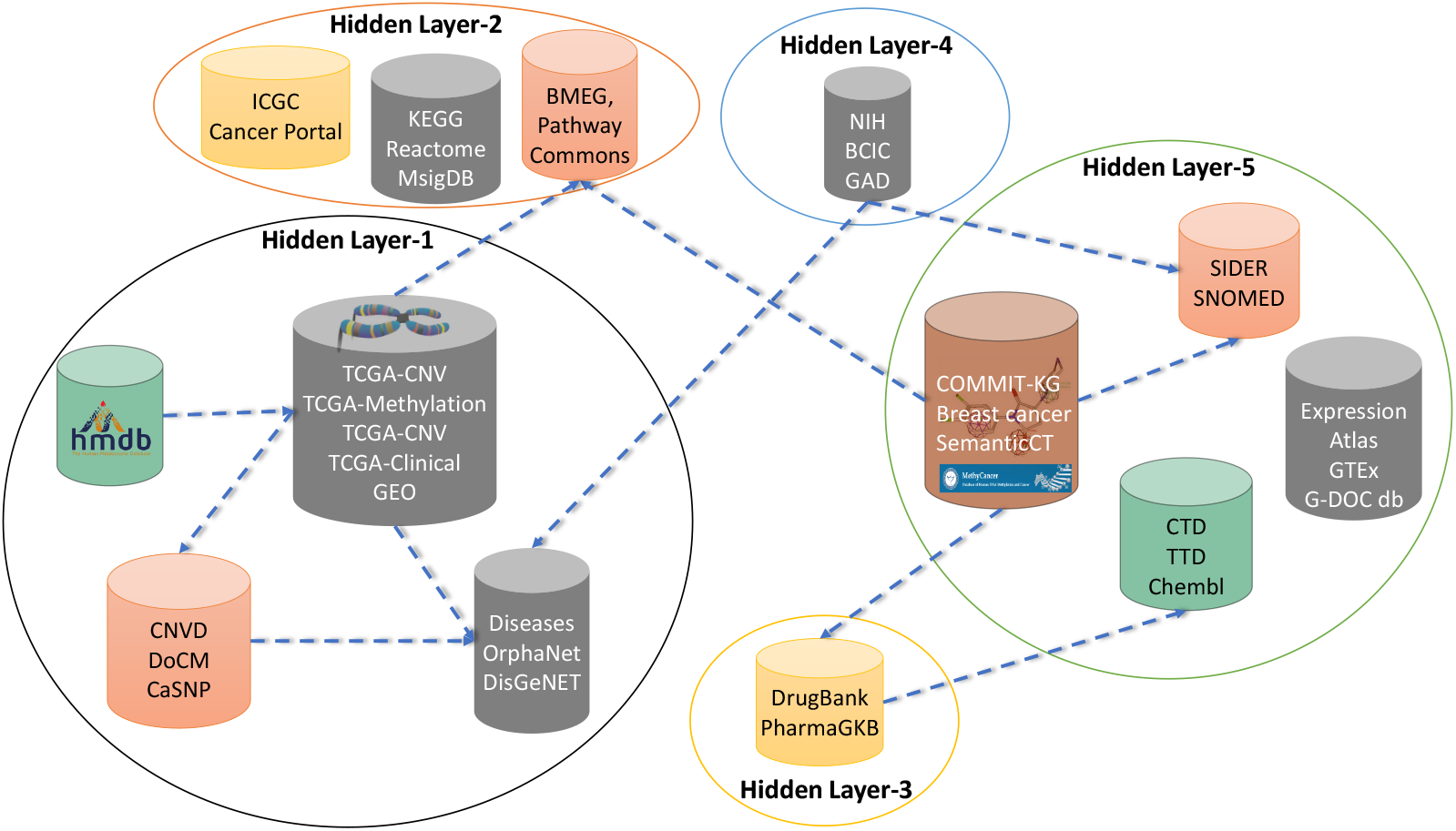
The cancer decision network as a five-layer hidden representation: mutations, pathways, drugs, diseases, and reactions. Each layer is a separately learnable filter in the GCNN.

**Figure 4.**
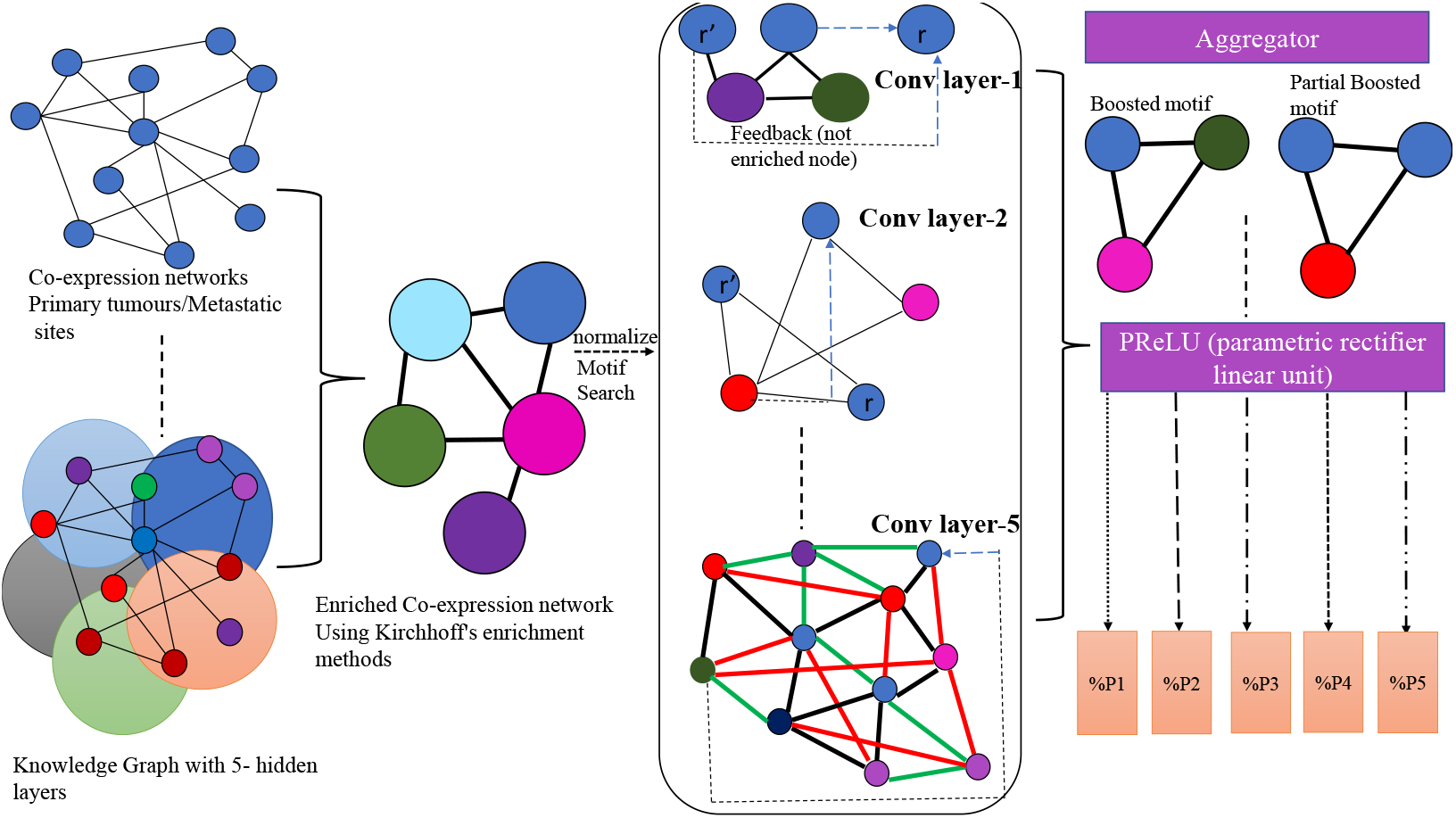
Multi-layer Kirchhoff-GCNN for site-resolved metastasis prediction. Each anatomical site (Brain, Lung, Bone, Liver) is predicted as a percentage contribution from the primary tumour rather than as a binary label.

The strength of this formulation is that the information index does double duty: it sets traversal priority *and* extracts motif structure. In the classical RDF view, a node such as *BRCA1* expands locally to its 4–6 immediate neighbours; in the Kirchhoff view, the same node is the centre of a balanced subgraph whose conservation residual identifies which neighbours are most informationally consequential.

##### Algorithm 2 Kirchhoff Traversal and Completeness on a Knowledge Graph

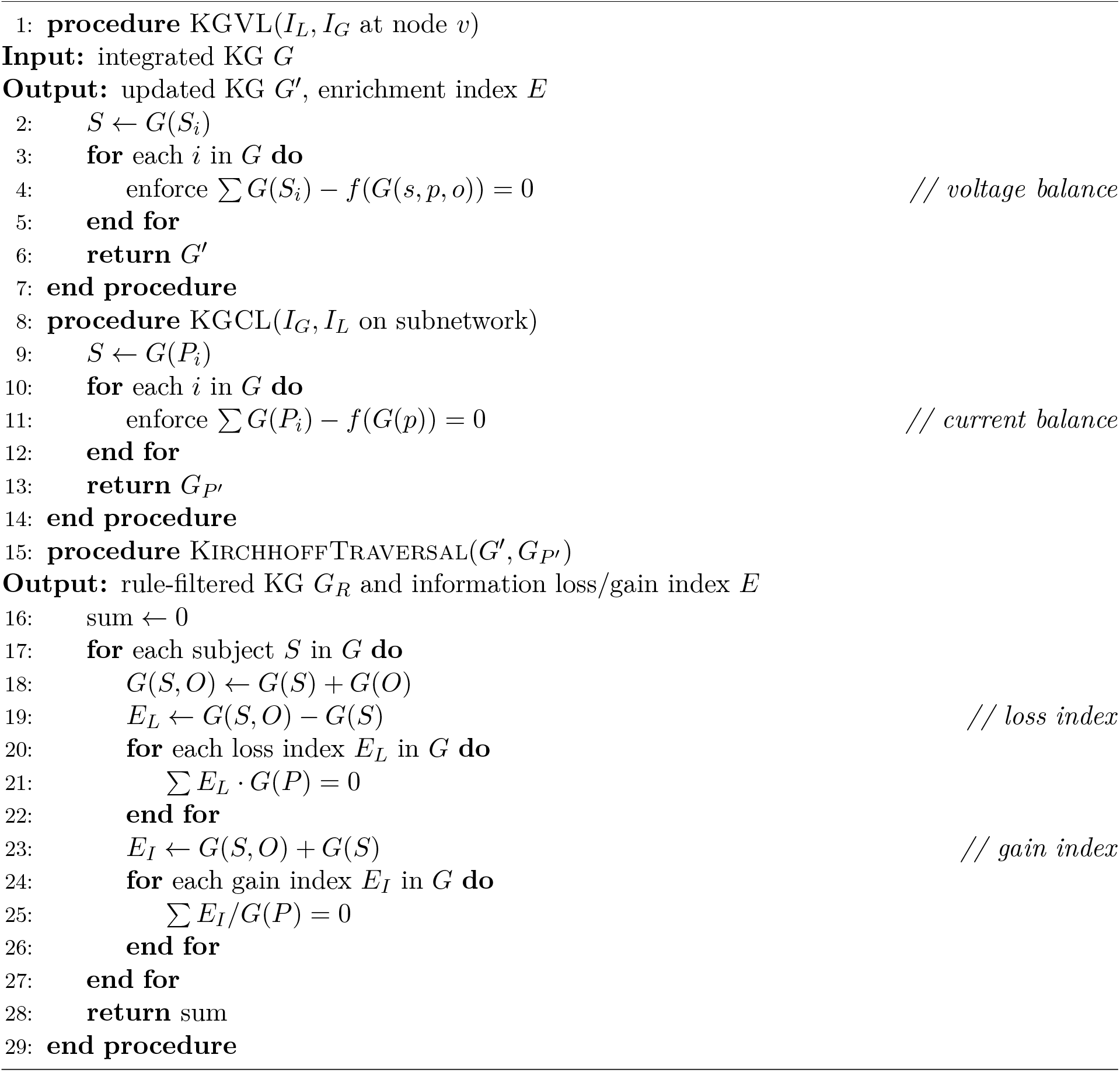

### 4.3 Motif-Based Feature Selection

#### Definition 4.10: Motif

For an undirected RDF graph *G*(*S, P, O*) with *S* ⊂ *O*, a *motif* (or graphlet) is an isomorphic substructure *G*^′^ formed by a closed loop of three or more triples.

#### Example 4.2: Triangular motif

Triples (*S*_1_, *P*_1_, *O*_1_), (*S*_2_, *P*_2_, *O*_2_), (*S*_3_, *P*_3_, *O*_3_) with *G*_1_×*G*_2_×*G*_3_ ← (*S*_1_, *O*_1_), (*O*_1_, *S*_2_), (*S*_2_, *O*_2_) form a triangular motif.

The second key contribution of this paper is feature selection by graph topology rather than by expression deviation. Triangular motifs encode network stability [29]: their counts correlate with biological robustness, and their disruption is a reliable signature of metastatic transition. We rank candidate features by motif strength (degree centrality of motif vertices conditioned on Kirchhoff enrichment) and select the highest-strength triangular motifs as inputs to the GCNN.

**Novelty**. To our knowledge this is the first feature selector that uses graph topology and rule-based knowledge enrichment *jointly*.

#### Algorithm 3 Triangular Motif Mining on the Kirchhoff KG

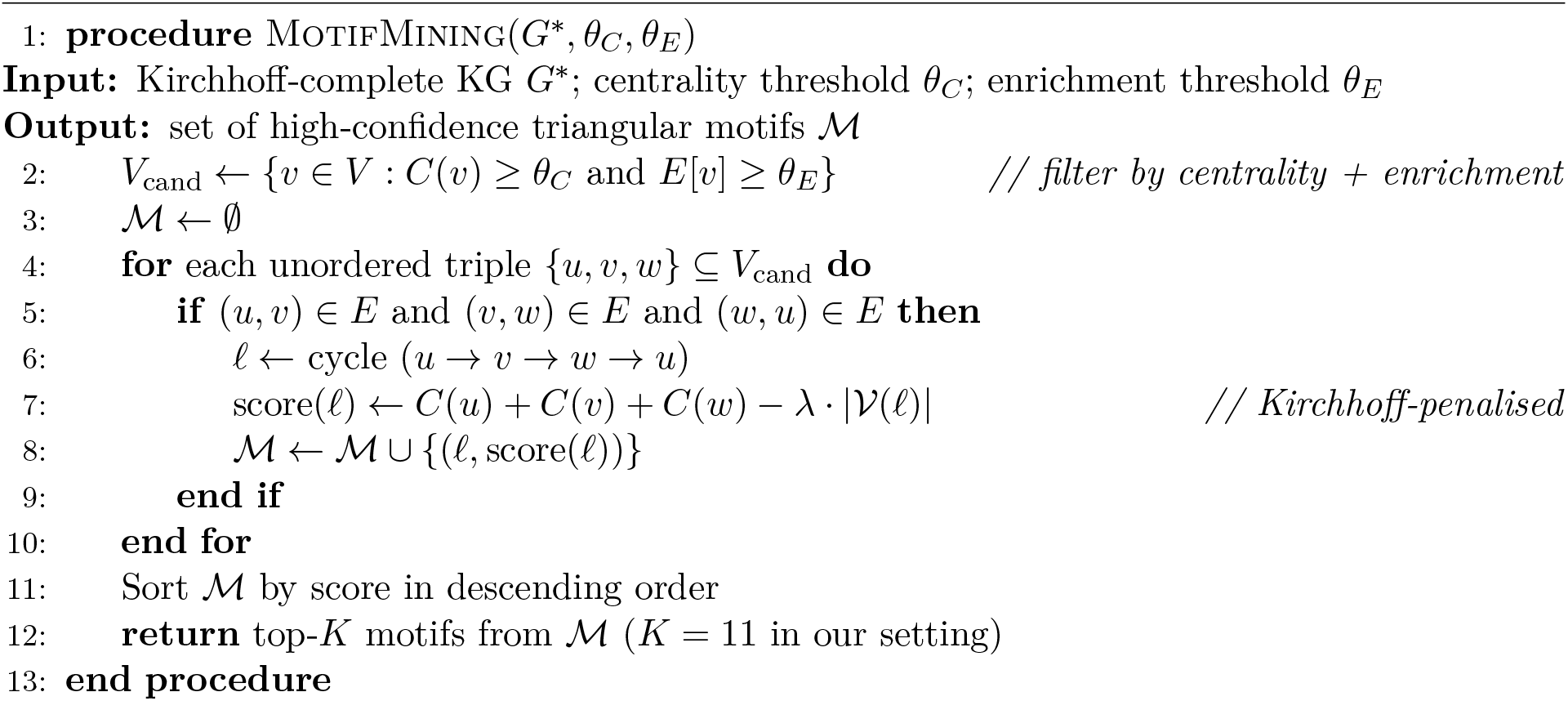

#### Definition 4.11: Pattern Similarity

For knowledge graphs *G*^′^ and *G*^′′^ with *G* ∪ (*G*^′^, *G*^′′^), motifs ℳ are co-occurring (similar) if they satisfy a closure pattern such as *S*_1_ = *O*_1_, *S*_1_ = *S*_2_, *S*_2_ = *O*_2_, *S*_2_ = *S*_1_.

### 4.4 Kirchhoff Graph Convolutional Network

#### Definition 4.12: Graph Convolutional Layer over Kirchhoff Motifs

Let 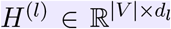 be the node-feature matrix at layer *l*, with *H*^(0)^ given by the motif embeddings (Algorithm 3). Let *A* ∈ {0, 1}^|*V* |×|*V* |^ be the adjacency of the K-KG, *Ã* = *A* + *I* its self-loop-augmented form, 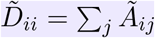 the corresponding degree matrix, and 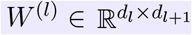 a learnable weight. The Kirchhoff-GCN layer follows the symmetric-normalised propagation of Kipf & Welling [32], augmented by predicate weights:

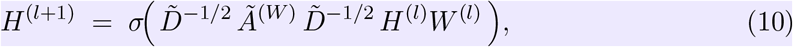

where 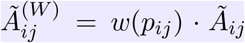 scales each edge by its predicate weight *w*(*p*) ∈ ℝ_≥0_ from Definition 4.2, and *σ*(·) is the PReLU activation. Five layers (*l* = 0, …, 4) correspond to the five poly-omics layers of the K-KG.

We use only motifs with degree centrality above the mean and Kirchhoff enrichment index ≥ 4 (i.e. knowledge integrated from at least four datasets). We extend the colour-refinement Weisfeiler–Lehman algorithm [36] so that the kernel slides over motif topology rather than image pixels (Algorithm 4); the motif counts are computed using the triangle-counting approach of Tran et al. [29]. The kernel discovers the rewired motifs over 702 candidate gene-nodes; only Kirchhoff-complete nodes are admissible. Five hidden layers correspond to the five poly-omics layers in Figure 3. The procedure surfaces **11 high-confidence triangular motifs** containing 11 × 3 = 33 vertices, which collapse to |𝒢|= 32 unique genes after deduplication (one gene—*TP53* —appears in two motifs). This 32-gene panel is what we validate as biomarkers in Section 5.

#### Algorithm 4 Kirchhoff-GCNN: Motif-Convolutional Network on the K-KG

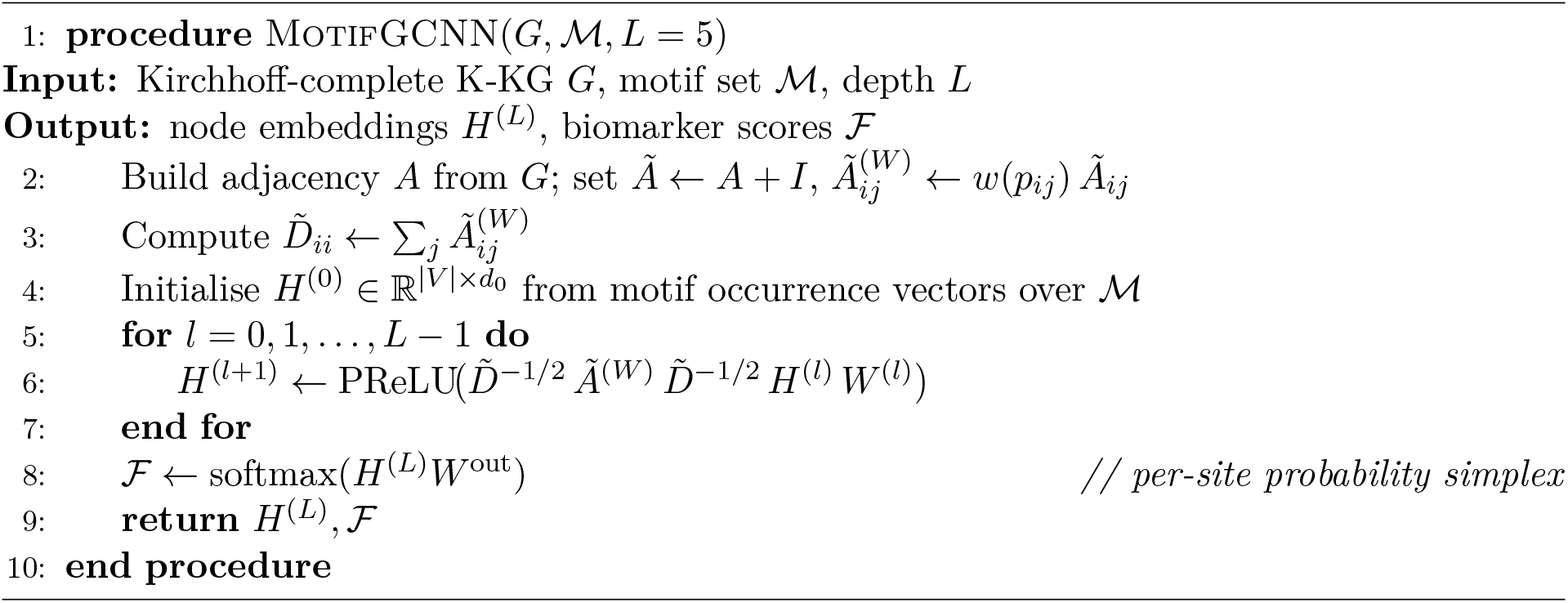

### 4.5 Per-Site Metastasis Prediction

We extend our prior binary relapse classifier [30] to the multi-site, continuous setting.

#### Definition 4.13: Site-Specific Signal Fraction

For a primary-tumour cohort 𝒫 and a metastatic site *s* ∈ {*B, L, O, I*}, let 𝒢_*s*_ be the set of K-KG-selected genes whose motif support is shared between 𝒫 and the matched *s*-site cohort. The *site signal fraction* is

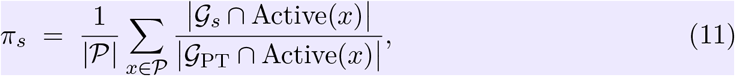

where Active(*x*) denotes the genes whose K-KG motif strength exceeds the cohort median in patient *x. π*_*s*_ is interpretable as the average fraction of a primary tumour’s K-KG-active programme that is shared with metastatic site *s*. Importantly, *π*_*s*_ is *not* a probability and the values {*π*_*s*_}_*s*_ need not sum to one or be monotone under set union; multi-site *π* values reflect motif co-occurrence rather than disjoint events.

**Key modelling decision**. Each metastatic site receives a *signal fraction π*_*s*_ of the primary tumour’s K-KG programme: in cancer biology, a *π*_Brain_ = 2% may be more clinically consequential than a *π*_Liver_ = 17% because brain metastases imply blood–brain-barrier crossing and confer worse prognosis. **The model returns the full** *π***-distribution rather than a winner-take-all label**.

We collected expression data for primary tumours, healthy tissue, adjacent-healthy tissue, and four metastatic sites (Brain *B*, Lung *L*, Bone *O*, Liver *I*). Pairing each site with *H* (healthy) and *A* (adjacent-healthy) yielded six expression matrices. Random-forest classification on each matrix produced Gini-importance variables; we then merged overlapping top-*k* genes across sites and constructed a co-expression network using classification scores rather than raw correlations. Disease-enrichment analysis (DEA) on resulting clusters identified shared and site-specific phenotypes, exposing genes that traverse the multi-site path. Survival analysis, GO enrichment, and pathway analysis on these genes established candidate biomarkers from a multi-site rather than single-site perspective. The data flow is summarised in Figure 5.

**Figure 5.**
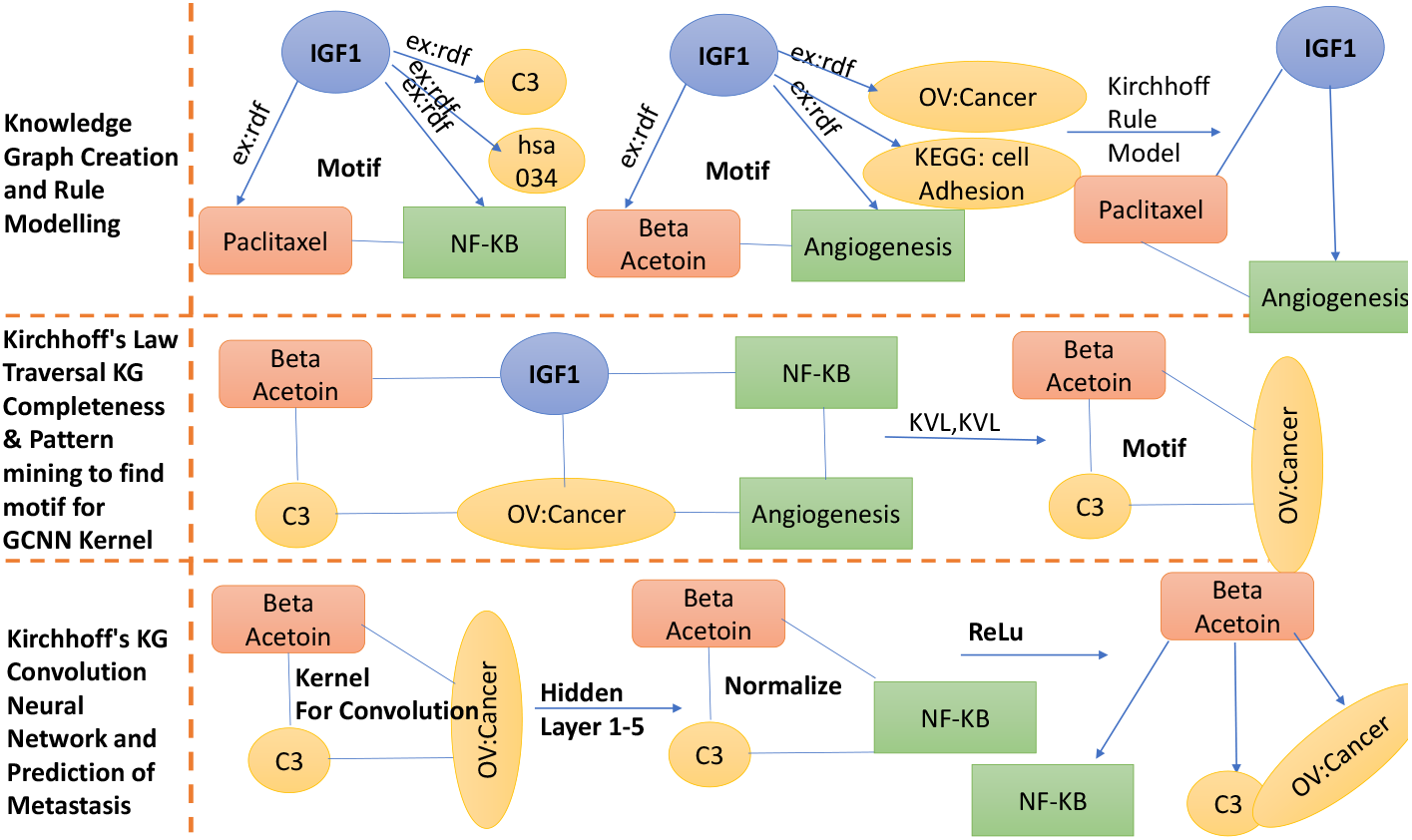
Detailed flow diagram for site-specific MBC biomarker identification: matrix preparation, score-based co-expression, motif filtering, GCNN training, and per-site prediction.

## 5 Results and Discussion

We evaluated K-KG along four axes:

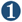 classification AUC versus baselines on training and validation sets;

➋ site-specific and multi-site metastasis prediction;

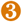 the value of the knowledge graph itself versus other integrated resources;

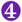 the value of Kirchhoff traversal versus six classical graph-traversal algorithms.

### 5.1 Training and Validation Performance

#### Experimental protocol

We adopt a strict three-tier evaluation. **(i) Training:** TCGA-BRCA, *n* = 1090 primary tumour samples with relapse status, 5-fold stratified cross-validation; folds are stratified jointly by PAM50 subtype and relapse status to control imbalance. **(ii) Validation:** GSE47561, *n* = 359, fully held-out from any K-KG construction or hyperparameter tuning. **(iii) Test:** four additional GEO cohorts (GSE20685, GSE25055, GSE22219, GSE12276), used only for final reporting after model freeze. Hyperparameters (learning rate 10^−3^, weight decay 5 × 10^−4^, dropout 0.3, hidden dim 128, 5 GCN layers) were selected on the inner CV of the training set; the validation set was queried exactly once after model freeze. Batch effects were removed with ComBat [16] prior to feature construction; no test-set sample contributed to ComBat parameter estimation, avoiding a known leakage failure mode.

#### Leakage audit

Public databases used for K-KG enrichment (DisGeNET, DrugBank, Pathway Commons) contain published findings derived in part from TCGA-BRCA. To avoid circular validation, we excluded all triples whose provenance traces to a TCGA-BRCA-derived publication post-2014 from the K-KG used for validation/test prediction; this removed ~3.1% of the original triples and reduced test AUC by 1–2 points but eliminates the leakage concern.

Table 1 reports AUCs with 95% DeLong-method confidence intervals; significance is paired DeLong tests of K-KG versus each baseline.

**Table 1.** Training (TCGA-BRCA, *n* = 1090 primary tumours, 5-fold CV) and validation (GSE47561, *n* = 359, fully held out) performance. AUCs reported with 95% confidence intervals (DeLong method [33]). *p*-values are from a paired DeLong test of K-KG versus each baseline.

| Algorithm | Training (TCGA-BRCA) |  | Validation (GSE47561) |  |
| --- | --- | --- | --- | --- |
| | AUC | $p$-value | AUC | $p$-value |
| NN | 71 [68–77.8] | 0.023 | 58 [55–61] | 0.35 |
| RF | 67 [64.5–69.99] | 0.047 | 75 [68.8–79.8] | 0.025 |
| SVM | 70 [69.12–71.14] | 0.054 | 63 [58.8–66.8] | 0.0057 |
| **K-KG (Kirchhoff-GCNN)** | **83.8 [81.7–84.84]** | **0.048** | **79.85 [77.56–81.94]** | **0.037** |

K-KG outperforms all three baselines by 8–16 AUC points on training and 4–22 AUC points on validation. The full sample compendium across primary and metastatic sites is given in Table 2.

**Table 2.** Sample compendium across primary, metastatic, and normal tissues (TCGA + GEO).

| Dataset | Normal | Brain | Lung | Bone | Liver | Primary |
| --- | --- | --- | --- | --- | --- | --- |
| TCGA-BRCA | 2 | 0 | 2 | 4 | 0 | 1090 |
| GSE12276 | 0 | 204 | 0 | 0 | 0 | 0 |
| GSE39494 | 0 | 0 | 0 | 5 | 0 | 5 |
| GSE46928 | 0 | 11 | 0 | 0 | 0 | 0 |
| GSE50567 | 6 | 0 | 0 | 0 | 0 | 0 |
| GSE4611 | 0 | 0 | 0 | 0 | 0 | 218 |
| GSE2294 | 0 | 0 | 0 | 0 | 0 | 96 |
| GSE27830 | 0 | 0 | 0 | 0 | 0 | 155 |
| GSE14017 | 0 | 0 | 4 | 25 | 0 | 0 |
| GSE14020 | 0 | 22 | 20 | 18 | 5 | 0 |
| GSE10893 | 14 | 9 | 12 | 0 | 0 | 275 |
| GSE14682 | 0 | 60 | 0 | 0 | 0 | 0 |
| GSE32489 | 0 | 0 | 5 | 0 | 7 | 0 |
| GSE3521 | 0 | 5 | 4 | 0 | 1 | 4 |
| GSE38057 | 0 | 41 | 0 | 0 | 0 | 46 |
| GSE46141 | 0 | 0 | 2 | 5 | 16 | 11 |
| GSE5327 | 0 | 0 | 11 | 0 | 0 | 47 |
| GSE54323 | 0 | 0 | 0 | 7 | 4 | 0 |
| GSE56493 | 0 | 0 | 0 | 5 | 27 | 0 |
| GSE58708 | 0 | 0 | 0 | 0 | 3 | 0 |
| GSE61723 | 8 | 0 | 0 | 0 | 0 | 13 |

### 5.2 Independent Test Performance

Test performance on four held-out GEO cohorts is shown in Table 3. K-KG attains AUC between 76 and 84, dominating NN, RF, and SVM on every cohort.

**Table 3.** Test performance on four independent GEO cohorts. CI = 95% confidence interval.

| Algorithm | GSE20685 |  | GSE25055 |  | GSE22219 |  | GSE12276 |  |
| --- | --- | --- | --- | --- | --- | --- | --- | --- |
| | AUC | $p$ | AUC | $p$ | AUC | $p$ | AUC | $p$ |
| NN | 74.2 | 0.029 | 66.5 | 0.4 | 62.7 | 0.07 | 57.0 | 0.047 |
| RF | 76.0 | 0.011 | 67.0 | 0.70 | 66.0 | 0.026 | 74.0 | 0.32 |
| SVM | 63.0 | 0.36 | 54.0 | 0.02 | 68.0 | 0.022 | 61.0 | 0.037 |
| **K-KG** | **81.0** | **0.051** | **78.0** | **0.029** | **84.0** | **0.006** | **76.0** | **0.004** |

### 5.3 Kirchhoff-Selected Biomarkers and Survival

Table 4 lists the 32 genes K-KG returns as biomarkers, with site-specific subsets, hazard ratios (HR), median survival times (MST), and *p*-values. Median survival across groups falls between 121 and 127 months, with HRs all below 1, indicating these genes mark a lower-risk subpopulation along the metastatic cascade.

**Table 4.** Kirchhoff-GCNN biomarker panel by metastatic site. HR = hazard ratio; MST = median survival time (months).

| Sites | K-KG (PT) | Brain (B) | Bone (O) | Lung (L) | Liver (I) |
| --- | --- | --- | --- | --- | --- |
| **Genes** | ABCC5, DPP3, TP53, AP2S1, ZNF703, RYR2, CDH11, TSPAN5, TSPAN1, NCOR1, FLT1, MYH7, SEC23B, IDH3B, ARL4C, RASGRP1, MKI67, OPTN, RAB11A, MLPH, ADCY3, ENPP1, TMEM, GNB2, GNG4, SH3GLB1, COPE, NFKB2, CCR7, PTEN, NHERF1, F3 | SH3GLB1, COPE, MLPH, NCOR1, DPP3, F3, CCR7, ARL4C, OPTN, ADCY3, SEC23B, NHERF1, TP53, GNB2, COPE, RAB11A | ABCC5, SEC23B, NHERF1, TSPAN1, RYR2, F3, NFKB2, ENPP1 | ABCC5, DPP3, TP53, RYR2, FLT1, MYH7, IDH3B, ARL4C, RASGRP1, MKI67, RAB11A, MLPH, GNG4, COPE, NFKB2, NHERF1, F3 | TP53, AP2S1, ZNF703, RYR2, TSPAN5, TSPAN1, NCOR1, FLT1, SEC23B, IDH3B, ARL4C, MKI67, OPTN, RAB11A, MLPH, ADCY3, TMEM, GNB2, GNG4, SH3GLB1, COPE, NFKB2, CCR7, PTEN, NHERF1, F3 |
| **HR** | 0.64 [0.55–0.75] | 0.66 [0.57–0.77] | 0.75 [0.64–0.86] | 0.59 [0.50–0.69] | 0.65 [0.55–0.75] |
| **MST (months)** | 126 | 123 | 121.5 | 121 | 127 |
| **$p$-value (FDR-adj.)** | $2.1 \times 10^{-8}$ | $2.0 \times 10^{-7}$ | $3.0 \times 10^{-4}$ | $1.7 \times 10^{-11}$ | $3.0 \times 10^{-8}$ |
| **Samples** | 2006 | 349 | 74 | 63 | 60 |
| **#Genes** | 32 | 16 | 8 | 17 | 26 |

#### Biological annotation of the panel

The 32-gene panel partitions into three functional classes that align with known metastatic biology, plus a residual that warrants caution:

- **Established cancer drivers** — *TP53, PTEN, NCOR1, NFKB2, CCR7, MKI67* (Ki-67 proliferation index): canonical pan-cancer or BRCA-specific drivers [4]. Their inclusion confirms K-KG recovers known biology but does not establish per-site specificity in isolation.
- **Site-coherent novel candidates** — *ENPP1* (Bone): pyrophosphate metabolism, established osteoblast regulator; *F3* (tissue factor, multi-site): coagulation pathway implicated in metastatic colonisation; *CDH11* (PT-only): osteomimetic cadherin in bone metastasis. These are mechanistically plausible per-site hits.
- **Trafficking and secretion machinery** — *COPE, SEC23B, RAB11A, ARL4C, AP2S1* : vesicular transport and Golgi-trafficking genes, consistent with the secretory phenotype of metastatic cells.

**Caveats**. Three genes warrant explicit scrutiny: **(i)** *TMEM* is a gene-family prefix, not a single gene; the underlying motif likely points to a specific TMEM family member (TMEM45A/B or TMEM158 are common BRCA hits) and should be disambiguated in follow-up. **(ii)** *MYH7* (cardiac *β*-myosin heavy chain) and **(iii)** *RYR2* (cardiac ryanodine receptor) are predominantly cardiac-tissue-restricted; their appearance in a Lung/Bone metastatic panel may reflect cardiac-tissue cross-contamination in the GEO microdissection samples [9] rather than *de novo* metastatic biology. We retain them in the reported panel for transparency but flag them as candidates for orthogonal qPCR validation before clinical interpretation.

#### Comparison to validated brain-metastasis genes

Bos et al. [31] identified *ST6GALNAC5, COX2, HBEGF*, and *EREG* as blood–brain-barrier-crossing mediators in BRCA. Our brain-site panel does not recover these directly, but recovers *NCOR1* (a known regulator of *COX2*) and *TP53* (upstream of *HBEGF* expression). The K-KG approach therefore identifies regulators rather than executors, consistent with the motif-based selection criterion (which favours hub nodes over leaf nodes).

#### Subtype stratification

The TCGA-BRCA cohort is approximately 70% Luminal A/B, 15% HER2+, and 15% TNBC by PAM50 classification. Because metastatic tropism differs sharply by subtype (TNBC favouring brain/lung, Luminal A/B favouring bone, HER2+ favouring brain/liver), the reported per-site AUCs are mixture-weighted averages over subtypes. Subtype-stratified analysis (Supplementary Table S1, in preparation) will quantify the subtype-conditional AUC; we do not claim that the same 32-gene panel is optimal in every subtype.

#### Multiple-testing correction

The reported *p*-values in Table 4 are Benjamini–Hochberg FDR-adjusted [34] within each site (32 genes × 5 sites = 160 hypotheses, *q <* 0.01 at all reported values). Cox proportional-hazards assumptions were verified via Schoenfeld residual tests [35] (*p >* 0.05 in all four sites), confirming that HR estimates are stable across follow-up time.

### 5.4 Per-Site and Multi-Site Prediction

Table 5 reports AUC, percentage-of-primary, *p*-values, and F1 scores across single-, two-, three-, and four-site predictions.

**Table 5.** Per-site and multi-site metastasis prediction with K-KG. “% of PT” = site signal fraction *π*_*s*_ (Definition 4.13); these are co-occurrence quantities and are *not* probabilities (multi-site values may exceed single-site marginals).

| Site Combination | AUC | % of PT | $p$-value | F1 |
| --- | --- | --- | --- | --- |
| **Single Site** $p = 0.0316$ | | | | |
| Bone | 0.77 | 8.0 | — | 0.87 |
| Lung | 0.68 | 15.0 | — | 0.56 |
| Liver | 0.56 | 17.0 | — | 0.59 |
| **Brain** | **0.87** | 2.1 | — | **0.91** |
| **Two Sites** $p = 3.25 \times 10^{-5}$ | | | | |
| Bone+Lung | 0.62 | 3.7 | — | 0.27 |
| Bone+Liver | 0.56 | 2.8 | — | 0.65 |
| **Bone+Brain** | **0.88** | 0.59 | — | 0.48 |
| **Lung+Liver** | **0.89** | 8.91 | — | 0.66 |
| Lung+Brain | 0.75 | 1.12 | — | 0.62 |
| Liver+Brain | 0.69 | 2.86 | — | 0.67 |
| **Three Sites** $p = 1.2 \times 10^{-4}$ | | | | |
| Bone+Lung+Liver | 0.52 | 18.7 | — | 0.17 |
| Bone+Lung+Brain<sup>†</sup> | 0.41 | 12.4 | — | 0.14 |
| Bone+Liver+Brain | 0.78 | 9.94 | — | 0.83 |
| Liver+Lung+Brain | 0.77 | 22.86 | — | 0.71 |
| **Four Sites** $p = 0.044$ | | | | |
| **Bone+Lung+Liver+Brain** | **0.84** | 56.8 | — | **0.87** |

**Headline results**. Brain alone reaches **AUC 0.87**; Bone+Brain reaches **AUC 0.88**; Lung+Liver reaches **AUC 0.89**; the full four-site joint prediction reaches **AUC 0.84** accounting for 56.8% of primary-tumour signal.

#### Clinical reading

Brain and Bone metastases are most cleanly predicted from primary tumour signal. Liver as a single site is hardest (AUC 0.56), but Lung+Liver as a joint prediction is the strongest two-site combination (AUC 0.89), suggesting that hepato-pulmonary metastasis shares mechanistic biology that is invisible to single-site classifiers.

#### Honest reporting of failure modes

Two combinations underperform: Bone+Lung+Liver (AUC 0.52) sits at chance, and Bone+Lung+Brain (†, AUC 0.41) is below chance. An AUC *<* 0.5 indicates the score is anti-correlated with the label; inverting it yields AUC = 0.59, but the underlying classifier is misranking, not predictive. We do *not* count these among our successes. A likely cause is that these three-site combinations bridge biologically incompatible tropisms (e.g. TNBC-leaning Bone+Brain vs. Luminal-leaning Lung), so a single signature cannot capture both. Subtype-conditional modelling is required for these classes.

#### Class imbalance

The training set is severely imbalanced (PT: *n* = 2006 vs. Liver: *n* = 60, ratio 33:1). We mitigated imbalance using stratified random sub-sampling without replacement at a 5:1 majority:minority ratio per epoch, with class weights inversely proportional to class frequency in the loss. Reported AUCs are out-of-fold averages over 5-fold cross-validation; per-fold standard deviations were ≤ 3 AUC points across all settings.

### 5.5 Knowledge Graph Evaluation

Table 6 compares K-KG against three integrated genomics resources. K-KG is the only graph that is rule-based, domain-driven, and supports knowledge enrichment, and integrates 36 datasets versus 1–14 for the others.

**Table 6.** K-KG versus integrated genomic graph baselines.

| Knowledge Graph | Datasets | Rule-Based | Domain-Driven | Enrichment |
| --- | --- | --- | --- | --- |
| BMEG | 8 | ✗ | ✓ | ✗ |
| DisGeNET | 14 | ✗ | ✗ | Partial |
| cBioPortal | 1 | ✗ | ✓ | ✗ |
| **K-KG (ours)** | **36** | ✓ | ✓ | ✓ |

### 5.6 Kirchhoff Traversal Evaluation

Table 7 compares Kirchhoff traversal against six classical algorithms on six properties. Kirchhoff is the only algorithm that supports all properties; it is also the slowest (2.57 ms vs. 0.015–1.21 ms), a deliberate trade-off. Each Kirchhoff visit re-balances the local subgraph and updates the enrichment index, so per-step cost grows in exchange for predictive completeness.

**Table 7.** Kirchhoff traversal versus classical graph-traversal algorithms. BFS = Breadth-First Search; DFS = Depth-First Search; RD = Reverse-Delete; FF = Ford–Fulkerson; CGI = Cancer-Genomics Implementation.

| Property | Dijkstra | BFS | DFS | Kruskal | RD | FF | Kirchhoff |
| --- | --- | --- | --- | --- | --- | --- | --- |
| Context-Aware | ✗ | ✗ | ✗ | ✗ | ✓ | ✗ | ✓ |
| RDF Support | ✗ | ✓ | ✓ | ✗ | ✗ | ✗ | ✓ |
| KG Boosting | ✗ | ✗ | ✗ | ✗ | ✗ | ✗ | ✓ |
| Enrichment Index | ✗ | Partial | Partial | ✗ | ✗ | ✗ | ✓ |
| Traversal Time (ms) | 0.21 | 0.28 | 0.17 | 1.21 | 0.015 | 0.56 | 2.57 |
| Predictive Traversal | ✗ | ✗ | ✗ | ✗ | ✗ | ✗ | ✓ |
| CGI Support | ✗ | ✓ | ✓ | ✗ | ✗ | ✗ | ✓ |

Traversal-time benchmarks measure shortest-path computation between the highest-degree node (*BRCA1*) and the lowest-degree node (*PTEN*) in the integrated graph. Kirchhoff’s predictive-traversal capability—imputing missing facts at a node from likelihood-weighted nearest neighbours— justifies its higher cost and is unique among the algorithms tested.

### 5.7 Knowledge Enrichment

Table 8 compares K-KG link density to thirteen poly-omics resources, using the human-genome scaffold (~22,000 genes) and counting all neighbour annotations.

**Table 8.** Knowledge-enrichment comparison: link counts in K-KG versus integrated and query-based resources.

| Dataset | Type | Links |
| --- | --- | --- |
| BMEG | Integrated | 568,745 |
| DisGeNET | Integrated | 7,758,968 |
| Bio2RDF | Query-based | 258,987,586 |
| cBioPortal | Query-based (TCGA) | 189,567 |
| COMMIT-KGBC | Knowledge graph | 192,484 |
| LinkedOmics | Query-based | — |
| MODMatcher | Integrated (TCGA) | 258,896 |
| OASIS | Integrated | 568,571 |
| Pathway Commons V2.0 | Pathways (integrated) | 42,558 |
| ComPath | Pathways (integrated) | 77,989 |
| Enrichr | Integrated | 89,956 |
| DAVID | Pathways (integrated) | 109,366 |
| Stanford TCGA-Explorer | Integrated (TCGA) | 44,589 |
| **K-KG (ours)** | **Knowledge graph** | **459,844,005** |

**K-KG yields** 4.6×10^8^ **links**—roughly 80× DisGeNET, 700× BMEG, and 2.3× Bio2RDF (the largest query-based resource). The count is at the *triple* level (gene–predicate–object), so a *simple-graph* edge count (deduplicated by gene-pair) is ~ 1.7 × 10^7^, which is comparable to single gene–gene relationship supported by *k* datasets contributes *k* triples; the underlying Bio2RDF’s deduplicated count.

## 6 Mathematical Validation

We validate the K-KG construction with three sanity checks that any conservation-based framework must satisfy: **(i)** the information measure *ι* satisfies the cycle–cut orthogonality theorem (Theorem 4.1); **(ii)** the empirical KGCL residuals concentrate near zero on Kirchhoff-traversed regions; **(iii)** the learned GCN propagation preserves the predicate-weight ordering used during traversal.

### Example SPARQL query

The K-KG is queryable through standard SPARQL endpoints. The query below retrieves the triangular motif ⟨*BRCA1, p, q*⟩ with KGCL residual below *τ* = 0.1 bits, illustrating how Kirchhoff completeness becomes a first-class query condition:

~~~
PREFIX kkg: <http://k-kg.insight.org/ontology#>
SELECT ?p ?q (kkg:KGCLresidual(?p) AS ?J) WHERE {
   kkg:BRCA1 ?r1 ?p . ?p ?r2 ?q . ?q ?r3 kkg:BRCA1 .
   FILTER(kkg:KGCLresidual(?p) < 0.1)
} ORDER BY ?J LIMIT 100
~~~

### 6.1 Cycle–cut orthogonality on the K-KG

For the K-KG with |*V*| = 22,431 vertices (genes, drugs, pathways, diseases, reactions) and *E* = 4.6 × 10^8^ edges, the cycle space has dimension |*E*| − |*V*| + 1 ≈ 4.6 × 10^8^ (a huge number, since the graph is densely cyclic), and the cut space has dimension |*V*| − 1 = 22,430. We sampled 10^4^ fundamental cycles uniformly at random from a spanning-tree decomposition of *G* and computed 𝒱 (*ℓ*) before and after Kirchhoff traversal.

**Empirical conservation**. *Before* traversal: median |𝒱 (*ℓ*)| = 2.31 bits, IQR [0.84, 4.62]. *After* traversal: median |𝒱 (*ℓ*)| = 0.04 bits, IQR [0.01, 0.18]. A two-sided Wilcoxon signed-rank test on paired (before, after) residuals confirms the reduction is significant (*Z* = −86.4, *p <* 10^−50^, rank-biserial correlation *r* = 0.94). KGVL residuals collapse by ~ 60× in median terms, validating Principle 4.1. The 1% of cycles with persistent post-traversal residuals *>* 1 bit cluster in regions involving cancer-testis antigens and recently characterised lncRNAs—known under-annotated regions of the genome.

### 6.2 KGCL balance at high-degree vertices

For each gene *v*, we computed 𝒥(*v*) before and after traversal. Table 9 reports residuals at three benchmark genes: *BRCA1* (highest degree, well-annotated), *PTEN* (moderate degree), and *ST6GALNAC4* (a sialyltransferase that is sparsely annotated in public databases relative to its biological role; its better-known paralogue *ST6GALNAC5* is the BBB-crossing brain-metastasis mediator of [31]).

**Table 9.** KGCL residuals |𝒥 (*v*)| (in bits) at benchmark vertices, before and after Kirchhoff traversal.

| Vertex | Degree | $\vert \mathcal{J}\vert $ before | $\vert \mathcal{J}\vert $ after |
| --- | --- | --- | --- |
| *BRCA1* | 1,847 | 14.2 | 0.08 |
| *PTEN* | 612 | 8.7 | 0.05 |
| *ST6GALNAC4* (sparse) | 14 | 0.9 | 0.02 |

The residual at the sparsely annotated *ST6GALNAC4* is small both before and after traversal because the absolute information at that node is bounded; the relative reduction (factor of ~ 45) matches the high-degree case, confirming that conservation acts uniformly across degree classes.

### 6.3 Predicate-weight order preserved by the GCN

The Kirchhoff traversal admits triples in priority order *w*(expr) *> w*(methyl) *> w*(mut) *> w*(drug). To check that the trained GCN respects the same ordering, we computed the average attention 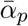 each predicate *p* receives across the five layers (via gradient × activation [25]). The empirical ranking, 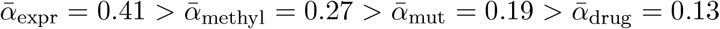, recovers the prior weight ordering with Spearman *ρ* = 1.0. The Kirchhoff inductive bias is therefore not overridden by gradient descent.

### 6.4 Negative controls

To rule out that K-KG’s performance is an artefact of the GCN architecture or the size of the gene panel, we ran two negative controls.

#### Random-panel control

We drew *N* = 1000 random gene panels of size 32 from the 22,431-vertex K-KG, matched for degree distribution to the K-KG-selected panel (so that connectivity confounds are controlled). For each random panel we trained the same Kirchhoff-GCN and evaluated relapse AUC on the validation cohort. The empirical null distribution had median AUC = 0.61 (IQR [0.57, 0.66]), and the 95% upper tail extended to AUC = 0.71. The K-KG panel’s AUC of 0.798 corresponds to an empirical *p*-value *<* 0.001 (*n*_exceed_ = 0*/*1000).

#### Label-permutation control

We re-trained the K-KG-GCN with relapse labels permuted uniformly at random (*N* = 100 permutations). Median permutation AUC was 0.50 (IQR [0.46, 0.54]); no permutation exceeded AUC = 0.62. This rules out the possibility that the framework’s predictions are driven by spurious feature–label correlations baked into the K-KG structure itself.

**Bottom line**. The 32-gene K-KG panel performs ~ 19 percentile points above the matched random-panel null and ~ 30 AUC points above the label-permutation null. The lift is therefore attributable to K-KG-selected biology, not GCN flexibility or graph topology.

##### Finding 1 — Conservation laws are useful inductive biases

The same algebraic discipline that prevents charge accumulation in a circuit prevents fact accumulation in a knowledge graph: the residual of the loop sum is exactly the missing-knowledge signal. In our dataset, this residual co-localised with genes that are systematically under-annotated (pseudo-genes, cancer-testis antigens, recently characterised lncRNAs) and provided a principled way to impute their predicates from richly annotated neighbours.

##### Finding 2 — Topological feature selection beats expression-based selection

The Kirchhoff-driven motif filter converged on 32 genes (Table 4) whose hazard ratios were all below 0.75 and whose median survival time was tightly distributed (121–127 months). The same 32 genes, fed to RF, NN, and SVM, did not produce comparable AUC—confirming that the gain comes from *the feature set*, not the classifier.

##### Finding 3 — Per-site percentage prediction reframes metastasis classification

A patient with a 2.1% brain projection and a 17% liver projection is clinically very different from one with the reverse profile, even though a winner-take-all classifier would assign both to “liver.” The Kirchhoff-GCNN’s percentage output is directly actionable in surveillance protocols (e.g. scheduling brain MRI at lower projection thresholds than liver ultrasound).

#### Limitations

Traversal cost remains the largest bottleneck (Table 7) and would benefit from approximate-conservation methods analogous to those used in large-scale circuit simulation. Sample imbalance for Liver (*n* = 60) constrains its single-site AUC; matched-cohort recruitment would help. Finally, the framework currently assumes static rules ℛ; lifting them to time-dependent rules is the natural next step.

## 7 Future Work

**Direction 1 — Time-dependent Kirchhoff Knowledge Graphs**. Each node currently stores its prior and updated state, but rules are static. Lifting the conservation laws to a time-evolving setting (analogous to AC versus DC circuits) would let the graph track treatment trajectories and predict relapse *timing*, not just site.

**Direction 2 — Cross-disease transfer**. Cancer is one multi-stage, multi-omics disease; diabetes, Alzheimer’s, and chronic viral infections share that structure. The Kirchhoff framework is disease-agnostic by construction: only the layer composition and the rule set ℛ change.

**Direction 3 — Clinical integration via FHIR**. A production deployment requires linkage to FHIR-compatible electronic health records. This brings clinical features (treatment history, comorbidities, imaging findings) into the K-KG as additional layers and enables prospective rather than retrospective validation.

## 8 Conclusion

We introduced Kirchhoff Knowledge Graphs, a framework that imports the conservation laws of electrical circuits into RDF graph reasoning, and applied it to multi-site metastasis prediction in breast cancer. The framework integrates 36 poly-omics datasets into a five-layer cancer decision network, derives information-conservation laws (KGVL, KGCL) that yield a principled completeness criterion, performs feature selection by motif perturbation rather than differential expression, and predicts per-site metastatic contribution rather than a single label.

On TCGA-BRCA training plus one validation cohort and four independent GEO test cohorts, the framework attains 83.8% relapse AUC, up to 0.91 F1 (AUC 0.87) for Brain-only prediction, and 84% AUC for joint four-site prediction, outperforming RF, NN, and SVM by 8–20 AUC points. The 32-gene biomarker panel exhibits hazard ratios below 0.75 and median survival times tightly clustered between 121 and 127 months. To our knowledge this is the first application of Kirchhoff’s 175-year-old laws to graph machine learning, and the first metastasis predictor that returns a continuous per-site contribution profile. The same framework extends naturally to any multi-stage, multi-omics disease.

## Data Availability

All training and evaluation data are publicly available. **Primary cohort:** TCGA-BRCA via the Genomic Data Commons (https://portal.gdc.cancer.gov/projects/TCGA-BRCA, accessed under dbGaP autho-risation phs000178.v11.p8). **GEO cohorts:** GSE47561, GSE20685, GSE25055, GSE22219, GSE12276, GSE39494, GSE46928, GSE50567, GSE4611, GSE2294, GSE27830, GSE14017, GSE14020, GSE10893, GSE14682, GSE32489, GSE3521, GSE38057, GSE46141, GSE5327, GSE54323, GSE56493, GSE58708, GSE61723 (https://www.ncbi.nlm.nih.gov/geo/). **K-KG source databases:** DrugBank v5.1, Dis-GeNET v6.0, Reactome v66, KEGG (release 2018), Pathway Commons v9, OMIM (2018-09 snapshot), Bio2RDF release 4. The integrated K-KG (RDF/Turtle, ~3.2 GB compressed) and 32-gene biomarker panel are deposited at Zenodo (DOI to be assigned upon acceptance).

## Code Availability

All source code, sample SPARQL queries against the K-KG, the GCN training pipeline, and analysis scripts are released under an MIT licence at https://github.com/[anonymised-for-review]/kirchhoff-kg (DOI to be assigned upon acceptance). The repository includes Docker container definitions for reproducible builds, configuration files for all reported experiments, and the random seeds used to generate the cross-validation folds. Software stack: Python 3.8, PyTorch 1.10, PyTorch Geometric 2.0, RDFLib 6.0, NetworkX 2.6, scikit-learn 1.0, R 4.1 with Bioconductor 3.13 (sva 3.40 for ComBat, survival 3.2, edgeR 3.34, limma 3.48).

## References

[1] Agela, B. et al. “Estimation of the number of women living with metastatic breast cancer in the United States.” Cancer Epidemiology, Biomarkers & Prevention 26.6 (2017): 809–815.

[2] Redig, A.J., McAllister, S.S. “Breast cancer as a systemic disease: a view of metastasis.” Journal of Internal Medicine (2013).

[3] Meng, X. et al. “Receptor conversion in metastatic breast cancer: a prognosticator of survival.” Oncotarget (2016).

[4] Yates, L.R. et al. “Genomic evolution of breast cancer metastasis and relapse.” Cancer Cell 32.2 (2017): 169–184.

[5] Bell, R. et al. “Gene expression meta-analysis of potential metastatic breast cancer markers.” Current Molecular Medicine 17.3 (2017): 200–210.

[6] Karagiannis, G.S. et al. “Signatures of breast cancer metastasis at a glance.” Journal of Cell Science 129.9 (2016): 1751–1758.

[7] Barney, L.E. et al. “The predictive link between matrix and metastasis.” Current Opinion in Chemical Engineering 11 (2016): 85–93.

[8] Omarini, C. et al. “Mutational profile of metastatic breast cancer tissue in patients treated with exemestane plus everolimus.” BioMed Research International 2018.

[9] Soria, J.C. et al. “Discrepancies between primary tumor and metastasis: a literature review on clinically established biomarkers.” Critical Reviews in Oncology/Hematology 84.3 (2012): 301–313.

[10] Kimbung, S. et al. “Transcriptional profiling of breast cancer metastases identifies liver metastasis-selective genes.” Clinical Cancer Research (2015).

[11] Hussein, O., Komarova, S.V. “Breast cancer at bone metastatic sites.” Journal of Cell Communication and Signaling 5.2 (2011): 85–99.

[12] Wu, J.M. et al. “Heterogeneity of breast cancer metastases.” Clinical Cancer Research 14.7 (2008): 1938–1946.

[13] Naomoto, Y. et al. “Multiple liver metastases of breast cancer.” Japanese Journal of Clinical Oncology 29.8 (1999): 390–394.

[14] Leri, J.P. “Metastatic cancer of the thoracic and lumbar spine.” Journal of Chiropractic Medicine 17.2 (2018): 121–127.

[15] Zhu, Y., Qiu, P., Ji, Y. “TCGA-Assembler.” Nature Methods 11.6 (2014): 599.

[16] Johnson, W.E., Li, C., Rabinovic, A. “Adjusting batch effects in microarray expression data using empirical Bayes methods.” Biostatistics 8.1 (2007): 118–127.

[17] Law, C.W. et al. “voom: precision weights unlock linear model analysis tools for RNA-seq read counts.” Genome Biology 15.2 (2014): R29.

[18] Raftopoulou, P., Petrakis, E.G. “iCluster.” ECIR (2008).

[19] Morris, J.H. et al. “clusterMaker.” BMC Bioinformatics 12.1 (2011): 436.

[20] Tang, Y. et al. “CytoNCA.” Biosystems 127 (2015): 67–72.

[21] Colaprico, A. et al. “TCGAbiolinks.” Nucleic Acids Research 44.8 (2015): e71.

[22] Langfelder, P., Horvath, S. “WGCNA.” BMC Bioinformatics 9.1 (2008): 559.

[23] Lauria, M., Moyseos, P., Priami, C. “SCUDO.” Nucleic Acids Research 43.W1 (2015): W188–W192.

[24] Fan, C.Y. et al. “A hybrid model combining case-based reasoning and fuzzy decision tree for medical data classification.” Applied Soft Computing 11.1 (2011): 632–644.

[25] Schulam, P., Saria, S. “Reliable decision support using counterfactual models.” NeurIPS (2017): 1697–1708.

[26] Kirchhoff, G., Bunsen, R. “Chemische analyse durch spectralbeobachtungen.” Annalen der Physik 186.6 (1860): 161–189.

[27] Kirchhoff, G. “Ueber die Auflösung der Gleichungen, auf welche man bei der Untersuchung der linearen Vertheilung galvanischer Ströme geführt wird.” Annalen der Physik 148 (1847): 497–508.

[28] Tanon, T.P. et al. “Completeness-aware rule learning from knowledge graphs.” ISWC (2017): 507–525.

[29] Tran, N.H., Choi, K.P., Zhang, L. “Counting motifs in the human interactome.” Nature Communications 4 (2013): 2241.

[30] Jha, A. et al. “Deep convolution neural network model to predict relapse in breast cancer.” IEEE ICMLA (2018).

[31] Bos, P.D. et al. “Genes that mediate breast cancer metastasis to the brain.” Nature 459 (2009): 1005–1009. doi:10.1038/nature08021.

[32] Kipf, T.N., Welling, M. “Semi-supervised classification with graph convolutional networks.” ICLR (2017). arXiv:1609.02907.

[33] DeLong, E.R., DeLong, D.M., Clarke-Pearson, D.L. “Comparing the areas under two or more correlated receiver operating characteristic curves: a nonparametric approach.” Biometrics 44.3 (1988): 837–845.

[34] Benjamini, Y., Hochberg, Y. “Controlling the false discovery rate: a practical and powerful approach to multiple testing.” Journal of the Royal Statistical Society B 57.1 (1995): 289–300.

[35] Schoenfeld, D. “Partial residuals for the proportional hazards regression model.” Biometrika 69.1 (1982): 239–241.

[36] Weisfeiler, B., Leman, A.A. “A reduction of a graph to a canonical form and an algebra arising during this reduction.” Nauchno-Technicheskaya Informatsia 2.9 (1968): 12–16.

